# A laboratory framework for ongoing optimisation of amplification based genomic surveillance programs

**DOI:** 10.1101/2023.07.17.549425

**Authors:** Connie Lam, Jessica Johnson-Mackinnon, Kerri Basile, Winkie Fong, Carl J.E. Suster, Mailie Gall, Jessica Agius, Shona Chandra, Jenny Draper, Elena Martinez, Alexander Drew, Qinning Wang, Sharon C Chen, Jen Kok, Dominic E Dwyer, Vitali Sintchenko, Rebecca J. Rockett

## Abstract

Constantly evolving viral populations affect the specificity of primers and quality of genomic surveillance. This study presents a framework for continuous optimisation of sequencing efficiency for public health surveillance based on the ongoing evolution of the COVID-19 pandemic. SARS-CoV-2 genomic clustering capacity based on three amplification based whole genome sequencing schemes was assessed using decreasing thresholds of genome coverage and measured against epidemiologically linked cases. Overall genome coverage depth and individual amplicon depth were used to calculate an amplification efficiency metric. Significant loss of genome coverage over time was documented which was recovered by optimisation of primer pooling or implementation of new primer sets. A minimum of 95% genome coverage was required to cluster 94% of epidemiologically defined SARS-CoV-2 transmission events. Clustering resolution fell to 70% when only 85% of genome coverage was achieved. The framework presented in this study can provide public health genomic surveillance programs a systematic process to ensure an agile and effective laboratory response during rapidly evolving viral outbreaks.

## Introduction

The near-real time generation and sharing of SARS-CoV-2 genomes has enabled unprecedented international surveillance of SARS-CoV-2 evolution [1]. SARS-CoV-2 whole genome sequencing (WGS) has been enabled by the use of multiplex polymerase chain reaction (PCR) amplification to selectively amplify SARS-CoV-2 from clinical specimens. Several tiling primer schemes were rapidly designed after the identification of the novel coronavirus causing COVID-19 disease in January 2020 [2][3-5]. These methods have been employed at scale during large outbreaks and increasing case numbers. At the start of the COVID-19 pandemic, our laboratory initially implemented a long-tiled amplicon WGS protocol [5] which produced high quality consensus genomes in a circulating viral population with low levels of mutational changes. However, amplification-based sequencing methods are sensitive to mutations or indels within primer sites, thus resulting in inconsistencies in amplification across the viral genome. The variability (and potential non-amplification) of SARS-CoV-2 genomic regions ultimately affects overall genome coverage required for in-depth genomic surveillance. Ongoing viral evolution, the emergence of multiple variants of concern (VOC) over the course of 2021, combined with increased case numbers in New South Wales (NSW), Australia, led to varying reductions in genome coverage and increasing difficulty in providing the level of genomic resolution established at the beginning of the COVID-19 pandemic. For example, the VOC Omicron has accumulated in excess of 53 mutations in comparison to the Wuhan-Hu-1 strain [6] (which the original ARTIC method was designed to survey) and continues to evolve. Over 30 of these mutations are concentrated in the spike receptor binding domain (RBD) within the coding region of the primary vaccine antigen [7]. The inability to amplify and sequence even a small part of the genome, particularly within the RBD, can have substantial effects on the integrity of genomic surveillance programs, including the loss of the ability to reliably detect emerging SARS-CoV-2 lineages [8-10]. Furthermore, rapid identification of punctuated or saltational evolution may be impacted, leading to unrecognised circulation of variants with potential increased transmissibility, virulence, or capacity to evade vaccine or infection induced immunity.

The importance of maintaining high sequencing quality to accurately reconstruct viral evolution has been recognised [11]. Ongoing decline in the efficiency of factors associated with amplification based WGS (e.g., primer specificity, multiplex cross reactivities) placed additional demands on genomics laboratories to maintain consistent levels of sequencing depth and genome coverage within increasingly diverse viral populations [11]. SARS-CoV-2 genomic surveillance has illuminated the need for ongoing assessment and optimisation of amplification based WGS to ensure the sustained quality of data and the agility and usefulness of surveillance programs. This study describes a framework for optimisation of genome sequencing efficiency based on our experience with SARS-CoV-2 surveillance during the COVID-19 pandemic. We quantified genome coverage loss across three years of the pandemic using a number of SARS-CoV-2 primer schemes. We established quality metrics for genome coverage and primer efficiency and applied them to trigger protocol reviews and modifications. The validation methodology to optimise multiple primer schemes to improve genome coverage and to sustain high-quality genome sequencing for public health surveillance is also outlined.

## Methods

### Retrospective review of SARS-CoV-2 genome coverage

SARS-CoV-2 genomes sequenced between April 2020 and May 2022 in NSW at the Microbial Genomics Reference Laboratory, Centre for Infectious Diseases and Microbiology Laboratory Services (CIDMLS), Institute for Clinical Pathology and Medical Research (ICPMR) - NSW Health Pathology, Australia were included in this study. Sequencing was performed as a part of the NSW Health program of integrated public health genomic surveillance which additionally collected data on epidemiological links between cases. Sequences were mapped to the SARS-CoV-2 reference genome (National Center for Biotechnology Information (NCBI) GenBank accession MN908947.3) and analysed to determine genome coverage. Only genomes with more than 500,000 reads mapped to the reference genome were included in the retrospective genome coverage review. The genomes were then categorized into VOC based on Pangolin lineage designation (Version 3.0.3) [12, 13].

### Representative viral cultures, RNA extraction and serial dilutions

Viral cultures representing seven distinct SARS-CoV-2 lineages causing COVID-19 outbreaks in NSW were selected. Each of these cultures were grown from clinical samples that were SARS-CoV-2 RNA was detected by reverse transcriptase real-time PCR (RT-PCR) as previously described [14]. Total RNA was extracted from 200μL of culture supernatant using either the RNeasy Mini Kit (Qiagen) according to manufacturer’s instructions or the MagNA Pure 96 instrument (Roche) with MagNA pure 96 DNA and Viral NA Small Volume Kit (Roche). cDNA was generated for all samples using LunaScript RT SuperMix Kit (New England BioLabs). Sufficient volume was prepared to perform serial dilutions (10^−2^ to 10 ^-7^) of the cDNA from each culture. RT-PCR targeting SARS-CoV-2 N-gene was then performed for each sample dilution to measure the SARS-CoV-2 load as previously described [15].

### Amplification based whole genome sequencing (WGS)

Three previously published amplification-based sequencing protocols, Long Amplicon[5], ARTIC (v3 and v4) [3] and Midnight (v1) [4], and a newly designed protocol, CIDM-PH oMicron (CIDoMi) (see methods below) were used. Long Amplicon and ARTIC methods were prepared as described previously [3, 5, 16].

Midnight v1 and CIDoMi primer sets were prepared following the SARS-CoV-2 1200-bp protocol [4]. Briefly, PCR master mixes were prepared for each pool of primers. Each PCR contained 12.5μL Q5 High Fidelity 2x Master Mix (New England Biolabs), 1.1μL of pooled primers mixes at 10μM (final concentration of each primer was ∼10-11pM), with 2.5μL of template and molecular grade water to a final reaction volume of 25μL. Cycling conditions were as follows: initial denaturation at 98°C for 30s, followed by 35 cycles of: 98°C for 15s, 65°C for 5min.

For all amplification protocols, PCR amplicons for pools 1 and 2 were combined for each sample and purified with a 1:1 ratio of AMPureXP beads (Beckman Coulter) and eluted in 30μL of sterile water. Purified products were quantified using Qubit™ 1x dsDNA HS Assay Kit (ThermoFisher Scientific) and diluted to 1ng for library preparation. Sequencing libraries were prepared using Nextera XT (Illumina) according to the manufacturer’s instructions. Libraries were then sequenced to generate 75-bp reads on either the Illumina iSeq or MiniSeq platforms.

### Bioinformatics analysis

Raw reads were quality controlled using an in-house pipeline [17]. Briefly demultiplexed reads were quality trimmed using Trimmomatic v0.36 (sliding window of 4, minimum read quality score of 20, leading/trailing quality of 5 and minimum length of 36 after trimming). Reads were mapped to the reference SARS-CoV-2 genome (NCBI GenBank accession MN908947.3) using BWA-mem version 0.7.17, with unmapped reads discarded. Variant calling and consensus genome generation was performed using iVar (v 1.2). Average read depth and genome coverage were calculated using Samtools coverage (v 1.15.1). Genomes with >98% coverage and >100x average read depth were considered high quality and included in further analysis. Read depth per amplicon was calculated using mosdepth (v 0.3.4)

### Determining SARS-CoV-2 amplification efficiency

The mean depth of coverage of the entire genome was determined using Samtools coverage (v 1.15.1). The median depth of each non-overlapping amplicon was calculated separately using mosdepth (v.0.3.4). A sequence read fraction (SRF) was used as a quantitative measure of amplicon efficiency. SRF was calculated for each amplicon of each isolate by dividing the median depth of coverage per amplicon by the average depth of coverage of the entire genome. Amplicons with median SRF>1.0 were considered efficient, SRF between 0.05-1.0 were classified as inefficient and amplicons with median SRF<0.05 were designated critically underperforming in the amplicon pool.

### Optimisation of whole genome sequencing (WGS) amplification

Underperforming amplicons were optimised by increasing primer concentrations as previously described by the COVID-19 Genomics United Kingdom (COG-UK) Consortium guidelines [18]. Critically underperforming amplicons (SRF <0.05) were identified for three of the WGS amplification methods (ARTIC, Midnight v1 and CIDoMi). The corresponding primer concentrations for these amplicons were increased relative to other primers within their primer pool. Overperforming amplicons (SRF >1.5) had corresponding primer concentrations reduced within the primer multiplex. Rebalanced primer concentrations for all WGS methods were then tested against the same panel of representative SARS-CoV-2 cultures dilutions. Amplicon SRFs were then calculated for each optimised method and compared to standard equimolar pooling.

### Primer design for CIDM-PH oMicron (CIDoMi), a new amplification based whole genome sequencing (WGS) schema

CIDoMi, a new amplification based WGS method, was designed using PrimalScheme (version 1.3.2) [19]. Eight SARS-CoV-2 genomes from samples collected between December 2021 and February 2022 were aligned and used as input into PrimalScheme (Supplementary Material). Genomes represented VOC Delta AY39.1.2 and AY39.1.3), VOC Omicron (BA.1, BA.1.1, BA.1.17 and BA.2) and Lineage B (NCBI genome accession MN908947.3) sequences. Primers were designed to produce 1200-bp amplicons using default parameters. Resulting primer sequences were evaluated for primer misfmatches against an alignment of 274 VOC Omicron genomes circulating in Australia (accessed on gisaid.org 22-03-2022, Supplemental Material).

### Determination of minimum level of genome coverage for genomic clustering and genotyping

A set of 159 VOC Delta genomes with >98% coverage and available supporting genomic and epidemiological links were curated from our routine SARS-CoV-2 genomic surveillance. SRF for each of the amplicons was generated using the methods described above, and amplicons were then ranked by SRF from least to highest amplification efficiency. Genome regions of the lowest SRF were masked from the whole genome alignment using bedtools (v 2.25.0) to exclude the least efficient 5%, 15% and 25% from the genomes. The masked genomes, equivalent to 75, 85 and 95% genome coverage were aligned to the reference genome using MAFFT (v 7.487). Each new set of consensus genomes were clustered using single nucleotide polymorphism (SNP) distances from snp-dists (v 0.6) and consensus trees were drawn for each coverage mask using MAFFT and IQTree (v 1.6.7) as previously described [20]. Comparisons between median SRF for equimolar and rebalanced protocols were performed using 2-tailed Mann-Whitney tests.

## Results

### Optimisation of primer efficiency of amplification based WGS protocols

Amplicon efficiency across three WGS amplification methods was tested and optimised using a set of viral cultures representing wild-type SARS-CoV-2 (Lineage A.2.2 (n=1)), as well as WHO-designated VOC Beta (Lineage B.1.35 (n=1)), VOC Delta (n=4) and VOC Omicron (BA.1 (n= 1) and BA.2 (n=1)). Across all variants the median concentration of culture extract dilutions ranged from 1.6×10^9^ copies/μl (cycle threshold (Ct)=24) to 1.6×10^2^ copies/μl (Ct=35) (Supplementary Figure 1).

A total of 13/24 culture dilutions produced >98% genome coverage using both equimolar and optimised ARTIC v4 primers. As expected, the average Ct value of complete genomes was lower (Ct 29.5, range 25.58-34.84) than for incomplete genomes (Ct 35.9, range 33.58 - 38.04). However, equimolar primers demonstrated a broader range of SRF efficiencies, with 15.2% (15/99) of amplicons critically underperforming (SRF <0.05). These included amplicons predominantly in the ORF1ab region (amplicons 1, 3, 5, 7, 9, 10, 11, 13, 14, 18, 32, 33, 50, and 60) and one that spanned the E and M genes (amplicon 88). Upon primer rebalancing only one ARTIC v4 amplicons had median SRF <0.05 (Figure 1A). This was amplicon 72 within the Spike coding region.

**Figure 1.**
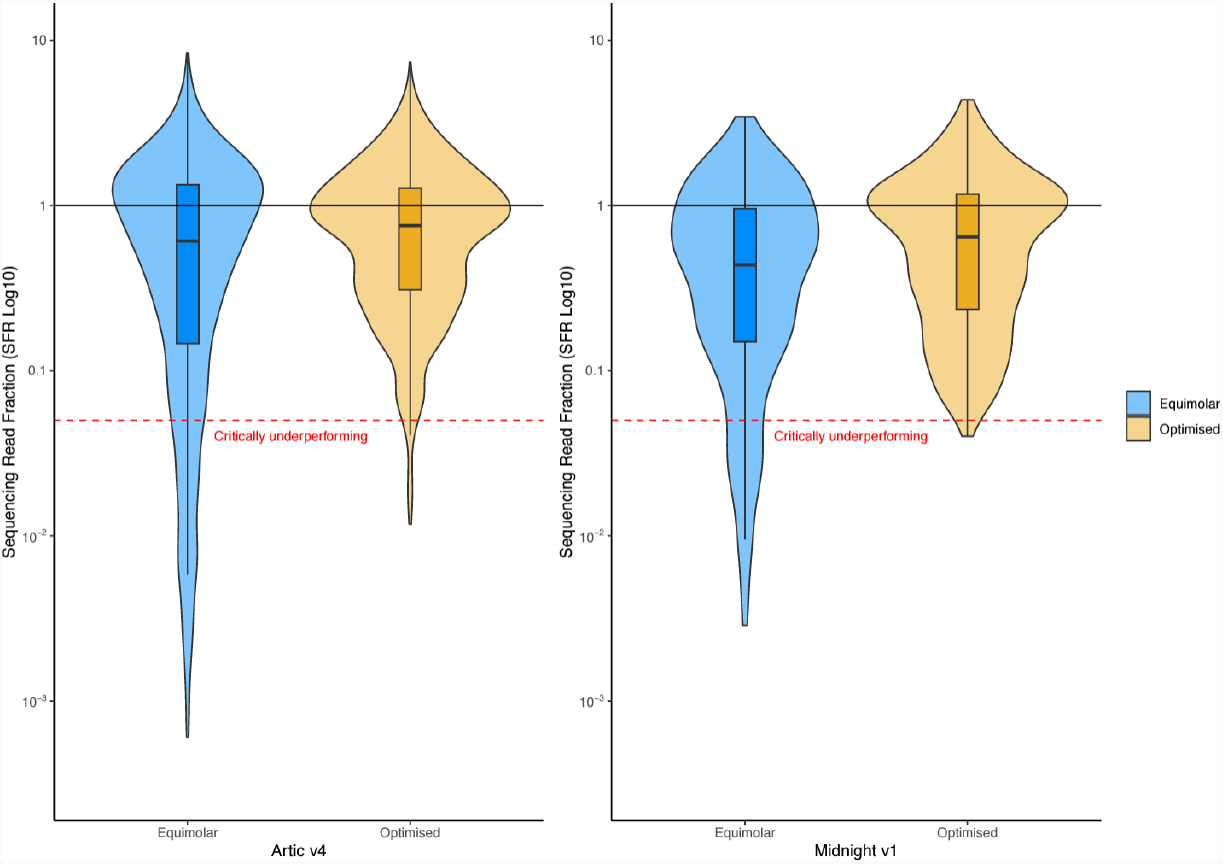
Violin and box plots for A) ARTIC v4 and B) Midnight v1 SARS-CoV-2 whole genome sequencing (WGS) methods. Blue violin and box plots represent equimolar primer pools whereas yellow violin and box plots have been calculated from primer pools which have been optimised and rebalanced based on sequence read fractions (SRF) of amplicons from the ARTIC v3 and Midnight v1 SARS-CoV-2 WGS protocols. The black line (SRF=1) represents ideal amplicon efficiency and the red dotted red line (i.e. SRF<0.05) is the threshold for critically underperforming amplicons. Medians for both optimised ARTIC v4 and Midnight v1WGS protocols are higher than their respective equimolar protocols, however no significant difference was observed (p>0.1).

WGS with Midnight v1 primers resulted in 54% (13/24) and 46% (11/24) of complete genomes from equimolar and rebalanced primers, respectively. Two dilutions failed library preparation after amplification with the optimised CIDoMi primers and were not repeated. The average SARS-CoV-2 Ct value of complete genomes was 29.55 (range 25.58-33.94) compared to 35.93 for incomplete genomes (range 32.85 - 38.22) (Supplemental Figure 1). However, 17.2% (5/29) of amplicons were critically underperforming using equimolar primers (SRF <0.05). This included two amplicons in the ORF 1ab region (2, 6), two amplicons in the spike gene (22 and 23) and one spanning the M, ORF6 and ORF7 genes (amplicon 27). After rebalancing primers of critically underperforming Midnight v1 amplicons all amplicons had a median SRF >0.05 (Figure 1B).

### Design and performance of CIDM-PH oMicron (CIDoMI) primers

SARS-CoV-2 VOC Omicron (lineage B.1.1.529) was first identified in South Africa in November 2021 but rapidly replaced VOC Delta as the dominant lineage throughout the world [21, 22]. A review of the SRF of 4,039 VOC Omicron genome sequences circulating in Australia during our study period indicated that three Midnight v1 amplicons (amplicons 10, 24, and 28) consistently failed to sufficiently amplify and produce adequate sequencing reads. In addition, 55% (16/29) of Midnight v1 amplicons had a median SRF<1.0 and three amplicons had SRF ≤0.05 (i.e., amplicons 10, 24, and 28). Further investigation identified mutations and indels within primer binding regions and suggested these amplicons could not be recovered by adjusting primer concentrations alone.

We addressed the loss of these informative regions by designing a new SARS-CoV-2 WGS protocol using Primal scheme [19] as described in the methods section. Eight Omicron VOC produced by our laboratory between December 2021 and February 2022 were used as representative Omicron sequences. Initial testing of CIDoMI with a dilution series of viral cultures of Omicron VOC BA.1 and BA.2 (Supplementary Figure 1) demonstrated that equimolar CIDoMI primers had a wide SRF distribution resulting in 5/11 genomes with >98% genome coverage. Although no individual amplicons had a median SRF <0.05, optimisation of CIDoMI primer concentrations increase the median SRF from 0.83 to 0.91 and generated >98% coverage from 9/11 isolate dilutions (Supplementary Figure 2).

### Setting thresholds for minimum genome coverage for variant detection and clustering

Genomic clustering was performed on a set of 159 SARS-CoV-2 VOC Delta (B.1.617.2) genomes and found to be consistent with reported transmission events and epidemiological context (Figure 2A). We estimated the effect of progressive loss of sequencing coverage on clustering capacity by masking critically underperforming amplicons (SRF<0.05) from the genome in a stepwise manner. Three levels of genome coverage loss were simulated, corresponding to the loss of 5% (5 amplicons), 15% (15 amplicons) and 25% (25 amplicons) of the genome. At 95% genome coverage, regions of the genome corresponding to five amplicons with the lowest SRF were excluded during clustering analysis (amplicons 1, 5, 9, 11, 18). At 85% genome coverage, genomic regions of an additional 10 amplicons (amplicons 7, 13, 10, 3, 60, 50, 33, 14, 23, and 88) were removed. The third simulation excluded a total of 25 amplicons to approximate 75% genome coverage (amplicons 29, 32, 64, 99, 44, 81, 26, 6, 77 and 42 in addition to the 15 amplicons mentioned above). At 95% genome coverage, resolution of genomic clustering was reduced but was still sufficient to correctly assign the majority (94%, 150/159) of isolates into their original clusters, or correctly identified as singletons (grey bars) (Figure 2B). Two isolates (1.3%, 2/159) which were previously in Cluster 1 could no longer be linked genomically, while five singletons (3%, 5/159) incorrectly joined Cluster 1. At 85% genome coverage (Figure 2C) the ability to distinguish Cluster 2 and Cluster 3 was lost as all 3 samples from Cluster 3, and 6 samples from Cluster 2 merged into Cluster 1; the remaining 2 samples from Cluster 2 became singletons. At 75% coverage the genomic clustering appeared markedly different from the original high coverage (>98%) clustering (Figure 2D). Only six clusters could be called with 75% genome coverage, compared with 11 clusters at >98% genome coverage. Furthermore, only 4 of these clusters (Clusters 8, 9, 10 and 11, Figure 2D) retained the same samples as the original 98% coverage clustering. The loss of genome coverage also led to increasing heterogeneity in Cluster 1 which expanded to include 12 isolates from other clusters and 14 singletons. A further six isolates could not be genomically assigned to a cluster and became singletons (3.8%, 6/159). Notably, none of the singletons clustered with each other to form “pseudo-clusters” at any of the genome coverage levels.

**Figure 2.**
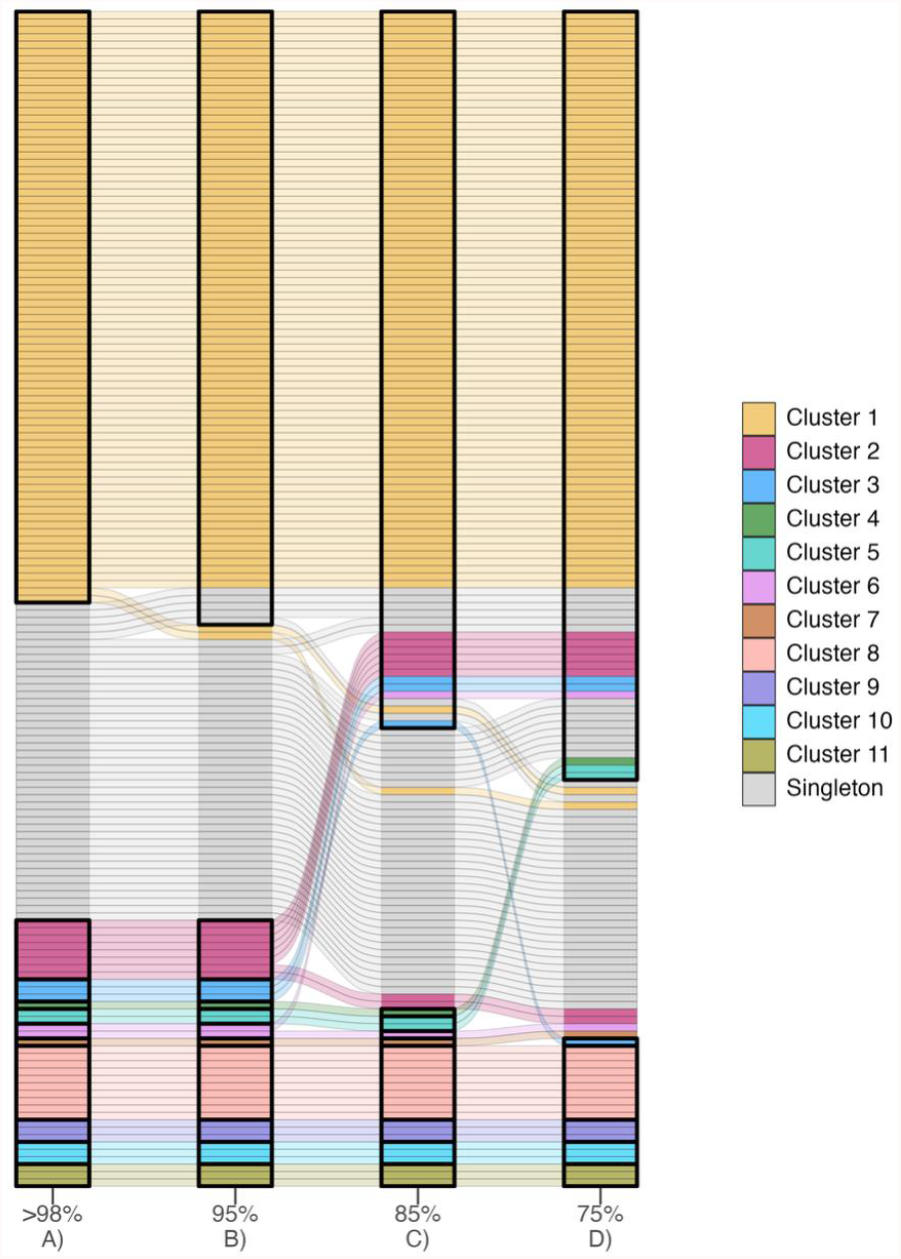
Changes to genomic clustering of 159 variant of concern (VOC) Delta samples as a result of simulated reduction in genome coverage. Each horizonal bar represents a single Delta genome which has been coloured according to its cluster membership in column A. Grey horizonal bars represent singleton Delta genomes which were not clustered. Vertical bold boxes outlined in black in column A) represent distinct epidemiologically confirmed genomic clusters, whereas boxes in columns B)-D) represent genomic clusters called when only 95%, 85%, and 75% of the genome is available for clustering analysis. The same set of 159 samples were used in each column. The lines between columns trace the movement of individual samples into different genomic clusters as the level of genome coverage is successively reduced.

### Monitoring genome sequencing coverage over time

We conducted a genome coverage review of SARS-CoV-2 genomes which were generated as part of public health genomic surveillance of cases diagnosed in NSW between February 2020 to May 2022. A total of 23,606 (98.8%) cases with >500,000 SARS-CoV-2 specific reads were included and were expected to generate complete genomes with a high depth of coverage. Between January 2020 and August 2020, average genome coverage was 98% using the long amplification protocol [5, 16] which gradually decreased to 95.7% in September 2020 (Figure 3B). ARTIC v3 was introduced in October 2020 to replace the long amplification protocol and to improve sensitivity of our genomic surveillance program as previously described. However, the ARTIC v3 introduction was associated with a further drop in genome coverage to 91.4% by November 2020. Following the widespread co-circulation of VOC Alpha and Beta in April 2021 genome coverage further declined to 75.1%. In October 2021, the Midnight v1 amplification protocol was implemented to better capture VOC Delta genomes and led to an increase in average genome coverage to 96.3%; however, subsequent changes in the viral population and the emergence of VOC Omicron (and associated sub-lineages BA.1 and BA.2) resulted again in a rapid reduction in genome coverage. Although the Midnight v1 amplification protocol performed well with Delta genomes, the average genome coverage for VOC Omicron strains between December 2021-April 2022 was only 83.6%.

**Figure 3.**
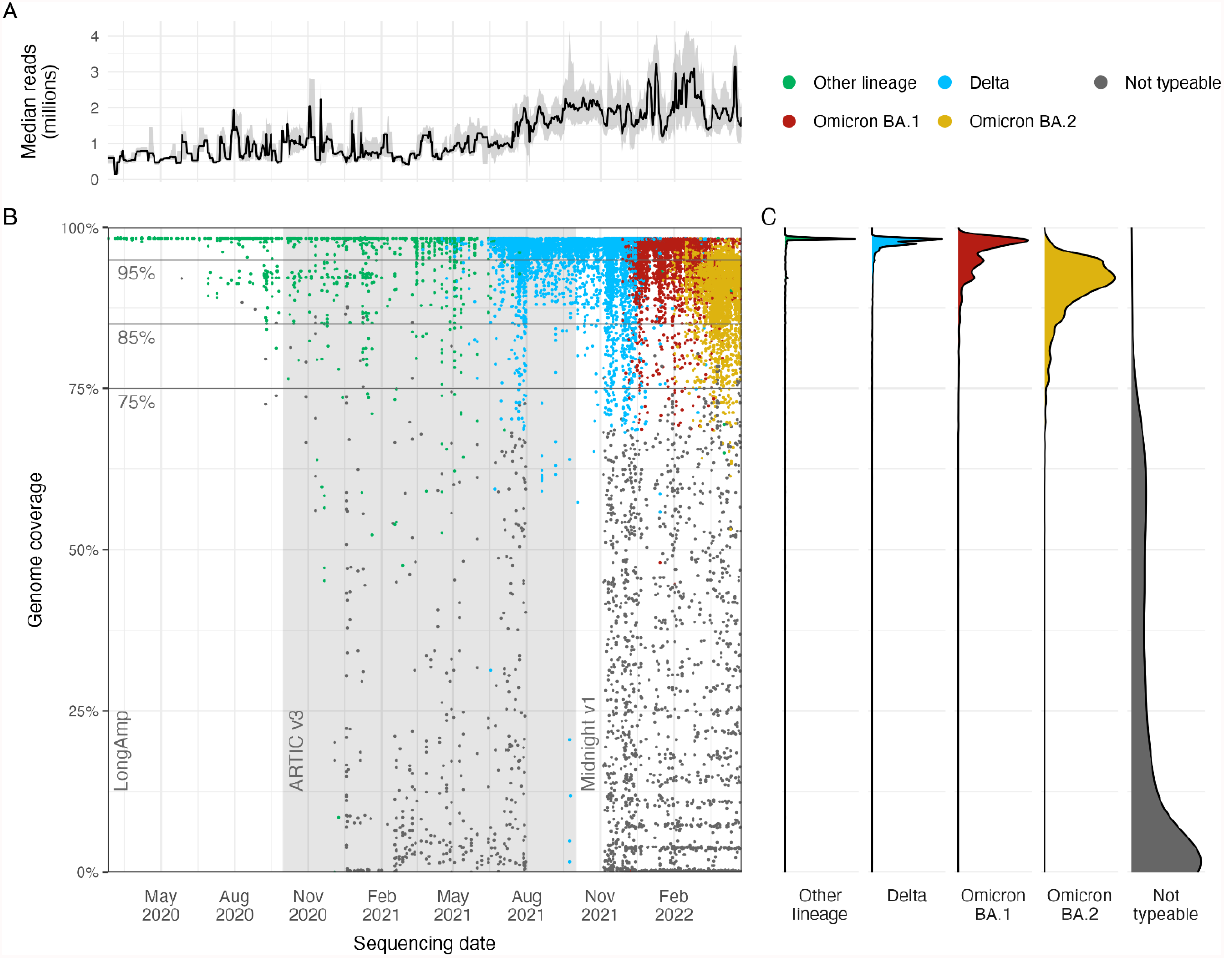
Genome coverage trends of 23,332 SARS-CoV-2 genomes collected in New South Wales (NSW) from January 2020 to the 31^st^ of April 2022. **A:** the upward trend in the median number of reads per genome over a rolling one-week period; interquartile ranges of reads are shaded in grey. **B:** Each coloured dot corresponds with a genome designated as variant of concern (VOC) Delta (blue), Omicron BA.1 (red) or Omicron BA.2 (yellow) or Not-typeable (grey). ‘Other lineages’ (green) include wild-type (Wuhan-like), Alpha, Beta and Gamma variants. Changes in sequencing protocols over time (Long Amplification, ARTIC v3 and Midnight v1) are labelled in the background of the plot, while the distribution of percent genome coverage for each VOC category is shown in panel C.

We compared the distribution of genome coverage of the VOCs Delta, Omicron, and non-typeable genomes to determine overall trends in sequencing efficiency per SARS-CoV-2 variant category (Figure 3C). The median genome coverage was 97.6% (IQR:92.6%-98.2%) Delta SARS-CoV-2, including wild-type, Alpha, Beta and Gamma VOCs (‘Other lineage’), 97.5% (IQR: 96.9%-98.3%) for Delta, 96.9% (93.9%-98.1%) for Omicron BA.1 and 91.7% (88.8%-98.7%) for Omicron BA.2, and 11.4% (0.3%-41.9%) for genomes that were not typeable.

A rolling 7-day median read count per genome is shown in Figure 3A. The increase in the number of reads produced per genome can be seen as a gradual trend as each successive VOC was introduced into NSW. During the initial phase of the COVID-19 pandemic when there was circulation of wild-type SARS-CoV-2, a median of 7.04×10^5^ reads were produced for each genome, which increased to 8.12×10^5^ and 1.00×10^6^ reads when VOCs Beta and Alpha began to co-circulate. The introduction of the VOC Delta and corresponding investment in high-depth sequencing resulted in a rapid increase to 1.78×10^6^ reads per Delta genome. A further increase, representing a tripling of the number of reads per genome compared to the start of the COVID-19 pandemic, was observed with Omicron BA.1 (2.11×10^6^ reads) and Omicron BA.2 (1.98×10^6^ reads).

### Sequencing efficiency optimisation framework for genomic surveillance

We established a framework (Figure 4) for monitoring and maintaining high quality genomes to provide adequate resolution for different levels of genomic surveillance. The 5-step framework was informed by ongoing optimisation of SARS-CoV-2 amplification based WGS methods employed at different times within our laboratory together with ongoing interrogation of the performance of these methods using routinely collected WGS metrics. Furthermore, the retrospective genome coverage review and simulation of genomic losses on surveillance resolution described in the results above, provide additional evidence to support the quality control parameters established.

**Figure 4:**
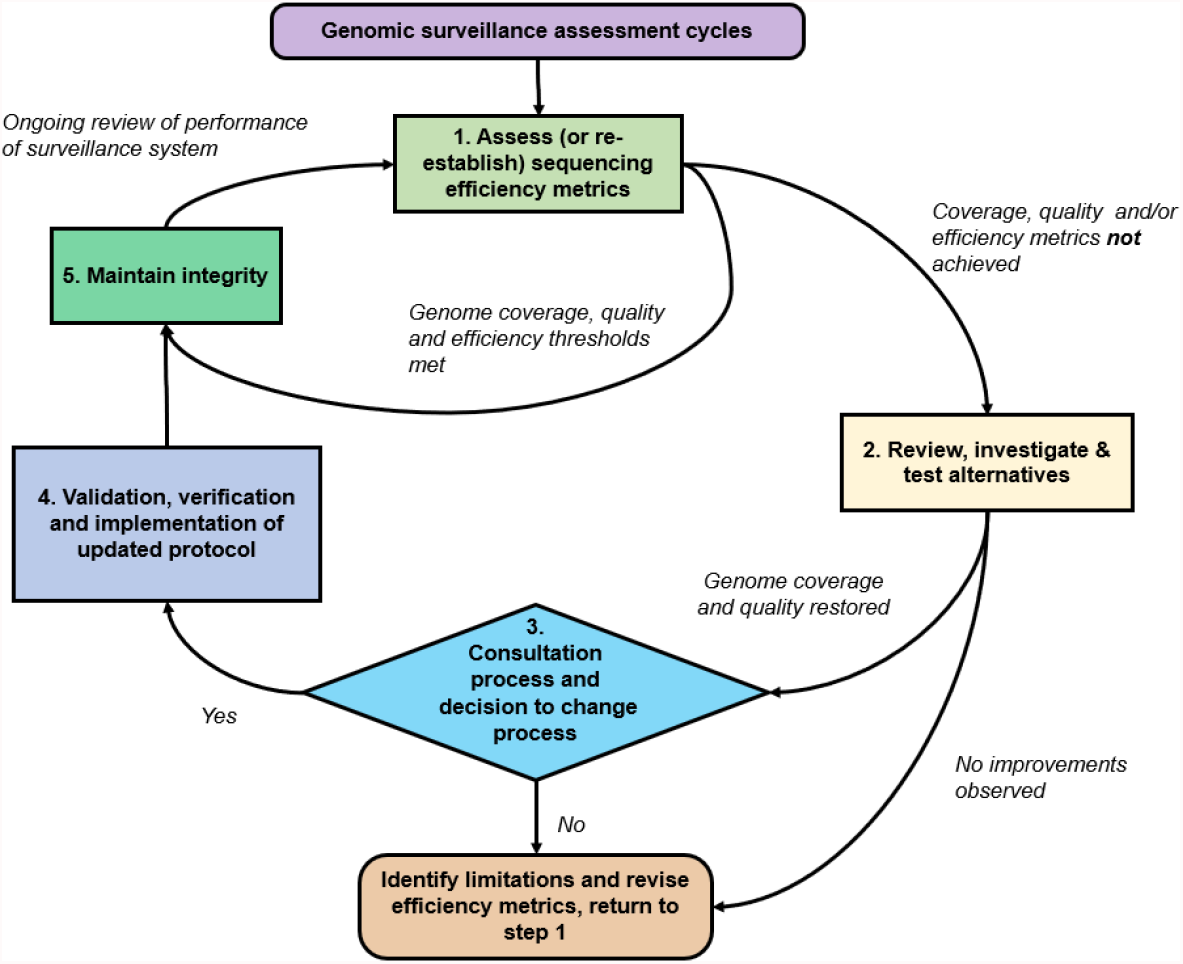
A quality framework to maintain and ensure high quality genomic surveillance of viral pathogens. The framework represents an ongoing iterative process to maintain a quality control within a genomic surveillance system, particularly when utilising amplification based whole genome sequencing (WGS). Each box represents an action that needs to be taken following the initial establishment of amplification based viral WGS, while the diamond indicates a decision that needs to be made after considering the outcomes of the WGS review, as well as any results from testing of alternative WGS methods. Directional arrows correlate with provide indications of observations of how the genomic surveillance system is performing.

The first step of the framework, after initial establishment of WGS surveillance is careful and ongoing monitoring of sequencing metrics collected during routine WGS. Using these metrics, appropriate benchmarks for genome coverage were set and reviewed periodically. Three levels of genome coverage (>98%, 95% and 85% genome coverage) were considered and corresponded to different levels of interpretation when used in genomic surveillance. When these benchmarks were not met, the second step of the framework was initiated. After successfully validating and implementing the ARTIC v3 protocol into routine SARS-CoV-2 WGS in October 2020, both gradual and sudden decreases in genome coverage below the lowest acceptable threshold of 85% were observed at different time points. Processes were then initiated to investigate reasons for poor sequencing performance (Figure 4, Step 2).

In November 2020, a substantial proportion of sequences (6.67%) were not able to be typed and by April 2021, average genome coverage had fallen to 75%, in large part due to the mutational changes in VOC Alpha and Beta. The review process revealed that the accumulation of mutations, particularly within the spike region, had resulted in the failure to amplify key regions of the SARS-CoV-2 genome. As this phenomenon was occurring at a global level, the ARTIC consortium released ARTIC v4 primers in June 2021 [23] which had been redesigned to accommodate high frequency mutations arising from VOC Alpha, Beta, Gamma and Delta. Other amplification based WGS protocols such as the Midnight protocol [4] had become available as alternative options. The 1200-bp tiled PCR protocol had been designed with the same principles as ARTIC, but with longer amplicons to accommodate for long-read sequencing technology, allowing it to fit into existing laboratory processes with negligible impact on sequencing workflow.

As part of the framework (Figure 4, Step 2), we proceeded to test, optimize and validate both methods against a selection of representative genomes (including the original wild-type SARS-CoV-2 (Lineage A.2.2), VOC Beta and Delta) to compare the performance of each method (Results described above). The results of each of the WGS methods, as well as other factors (including the ability of each method to perform at scale, amount of verification and validation, and potential disruption to existing laboratory workflows) were used in the decision-making process (Figure 4, Step 3) of whether to change existing workflows. In October 2021, routine SARS-CoV-2 WGS transitioned to using Midnight v1, and additional validation, verification processes were performed as part of the routine public health genomics surveillance service at our laboratory.

Each step of the framework was further refined when another iteration of decrease in genome coverage was observed. Substantial genomic changes within the viral population were again affecting the performance of the optimized sequencing protocol. Reports of sequences with a set of unusual mutations [24] and the first case of the SARS-CoV-2 lineage B.1.1.529 designated the VOC Omicron had been identified in NSW on November 27, 2021 [25]. While investigating alternative WGS methods or improvements to the Midnight v1 protocol, higher-depth sequencing was re-instated in order to sustain a high-resolution local tracking of cases infected with VOC Omicron. An update for ARTIC v4 was released in December 2021 involving the addition of 11 new primers [23], which also offered a potential to recover lost genome resolution. However, if implemented, the updated primers would still require optimization and rebalancing as well as updates due to the number of primers in the protocol and continuous representative surveillance. Given the characteristics and mutational profile of the VOC Omicron, we opted to design the CIDoMi protocol which provided us with over 99% of genome sequence in the circulating VOC Omicron genomes.

## Discussion

The integration of high-resolution genomic surveillance into the public health response in NSW enabled the following applications of SARS-CoV-2 genomic data: (i) near-real-time clustering of cases and uncovering cryptic transmission events to support outbreak response and public health investigations, (ii) lineage assignment, and (iii) detection and identification of new and emerging variants. Such integration of genomic surveillance into a pandemic response improved the understanding of local transmission dynamics and public health outcomes [20, 26]. This study describes cycles of continuous optimisation of sequencing efficiency which has been formalised in a quality framework (Figure 4), and has been essential for ensuring agility and sustainability of sufficient resolution for effective SARS-CoV-2 surveillance. Through iterative reviews, comparison of differences in genome coverage and SRF, we identified and investigated inefficient amplicons. Without optimisation, inefficient amplicons can cause uneven distribution of reads across the genomes; as a result, informative SNPs may not be captured in poorly amplifying regions, particularly when sequenced at lower depths.

The impact of viral divergence on amplicon sequencing performance was clearly demonstrated by a sharp reduction in genome sequencing coverage associated with the emergence of successive SARS-CoV-2 VOCs. The accumulation of genomic differences from wild-type SARS-CoV-2 to VOC Alpha, from Alpha to Delta, and particularly from Delta to Omicron was indicative of saltational evolution within the viral population [27] and likely driven by persistent infections within susceptible immunocompromised hosts [28]. Saltational events (i.e. large multi-mutational jumps resulting in distinct and distant variants) have been predominantly observed within the N-terminal domain and receptor binding domain of the spike protein and were also considered a feature of emergent SARS-CoV-2 VOCs [29]. An unexplained reduction in genome coverage and clustering sensitivity can be treated as a signal of significant viral evolution, impacting performance of sequencing protocols, once other explanations, such as laboratory contamination or index switching, are ruled out. Maintaining high quality genomic surveillance is thus crucial in detecting these events to provide accurate genomic information for the development of diagnostic assays targeting these regions.

Based on our findings, a minimum 95% genome coverage threshold was established as a requirement for high resolution genomic clustering of SARS-CoV-2, identification of cryptic transmission and disease transmission events, as well as early detection of variants. Reductions in clustering resolution were immediately apparent at a genome coverage of 95% but were still able to correctly identify the majority (94%) of epidemiologically linked clusters. Adequate quality genomes with coverage between 85-95% may still allow for genomic and epidemiological clustering if lineage defining SNPs reside within high performing amplicons. Lineage calling for these genomes may still also be feasible, which can then further guide the prioritisation of samples for enhanced investigation. Our curated set of VOC Delta genomes demonstrated that at 85% genome coverage genomic links between related isolates were unable to be made, and missing genomic markers contributed to incorrect clustering. Clustering of unrelated cases (singletons) or merging or misassignment of defined genomic clusters can have significant misleading effects on follow up public health actions. Our findings benchmarked the genome quality and coverage thresholds with a clear understanding of where incomplete genomic information can still be of value in a public health genomic surveillance system.

The decision-making process in this framework (Figure 4, Step 3) is a key step and should be informed by available alternative processes identified during the review and investigation stages of this framework (Figure 4, Step 2). Importantly, any decisions should only be made after consideration and consultation with relevant stakeholders who ultimately act upon the genomic data generated, including public health units and clinicians. For example, high genome resolution maybe required when investigating cases of treatment failure to inform the best antiviral treatment strategy, or for lineage designation of emergent variants. The benefits and limitations of changes to be made must also be conveyed during such assessment, as other key metrics may be affected, such as turnaround time for sequencing. The ability to produce and maintain high quality genomes for laboratory surveillance depends on a range of (external) factors, including the viral yield within a sample and the speed of emergence of new variants. Pre-analytical factors such as sample collection, transport and processing may interfere with the integrity of the original specimen and reduce the likelihood of recovering a complete genome [30]. While such external factors are difficult to control, internal factors, such as sequencing platforms and equipment, competencies of laboratory and bioinformatic personnel, can be modified within an organisation and are crucial factors in maintaining the integrity of the genomic surveillance system (Figure 4, Step 5).

In conclusion, the SARS-CoV-2 genome coverage and SRF can serve as informative indicators of reduction in performance due to viral evolution and act as triggers for sequencing protocol modifications. The proposed framework can help laboratories to sustain high-resolution amplification-based public health genomic surveillance through prolonged epidemics of rapidly evolving pathogens and evolving public health containment approaches. In the context of increasing demand and associated costs of generating pathogen genomes, this sequencing optimisation approach ensures the ongoing utility and accuracy of viral genomic surveillance programs. The COVID-19 pandemic informed the development of this framework which can be further adapted to be pathogen agnostic and universally applied to other viral pathogens with epidemic potential.

## Acknowledgements

The authors acknowledge the Sydney Informatics Hub and the use of the University of Sydney’s high performance computing cluster, Artemis. The authors are grateful to NSW Health Pathology partner laboratories, for referring samples for genomic surveillance. Expert advice and epidemiological information provided by the NSW public health surveillance units at the NSW Health Protection are also gratefully acknowledged. RR is supported by an Investigator Grant (GNT2018222) from the National Health and Medical Research Council, Australia (NHMRC).

## Author contributions

Study concept and design by CL, JJM, RR, VS. Sample processing and testing by CL, JA, JJM, RR, SC. Sequencing and analysis by CL, CS, KB, JJM, MG, RR, WF. The first manuscript was written by CL, KB, JJM, RR with additional editing from CS, JA, SC, WF, VS. The final manuscript was approved by all authors.

## Supplementary Data and Figures

### Design of new SARS-CoV-2 whole genome sequencing protocol CIDoMI

Sequence data for eight SARS-CoV-2 genomes used to design CIDoMI SARS-CoV-2 whole genome sequencing protocol is available here:

Eight SARS-CoV-2 genomes from samples collected between December 2021 and February 2022 were aligned and used as input into PrimalScheme (Supplementary Material). The eight genomes represented SARS-CoV-2 Lineage B (NCBI genome accession MN908947.3) and the following variants of concern: Delta AY39.1.2, Delta AY39.1.3, Omicron BA.1, BA.1.1, BA.1.17 and BA.2.

### Retrospective review of SARS-CoV-2 sequences

A total of 23,606 unique sequences were used for the retrospective review of SARS-CoV-2 whole genome sequencing between February 2020 and May 2022. Of these, 17,396 sequences met the upload criteria to GISAID and the sequence data for these samples are available in the GISAID EPI_SET**: <u>10.55876/gis8.230502km</u>**

Sequences which did not meet quality control criteria, duplicate sequences and non-typeable sequences have been excluded from the GISAID EPI_SET.

**Supplementary Figure 1.**
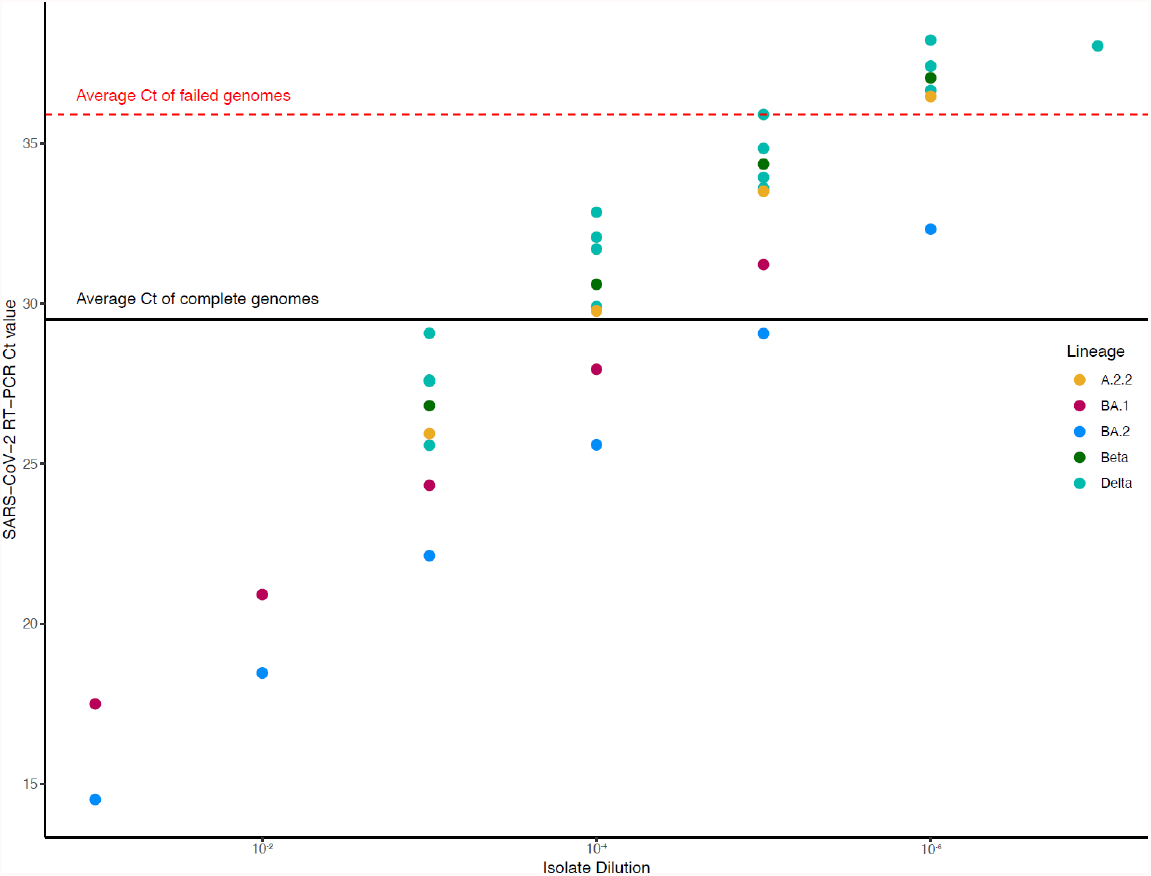
Real-time PCR cycle threshold (Ct) values of culture extract dilutions in this study. Serial dilutions of SARS-CoV-2 cultures corresponding to Lineage A.2.2.2 (yellow), Variants of Concern Beta (green), Delta (teal), Omicron BA.1 (pink), BA.2 (blue). The average SARS-CoV-2 Ct value of all complete genomes was 29.55 (range 25.58-33.94) compared to 35.93 for incomplete genomes (range 32.85 - 38.22).

**Supplementary Figure 2.**
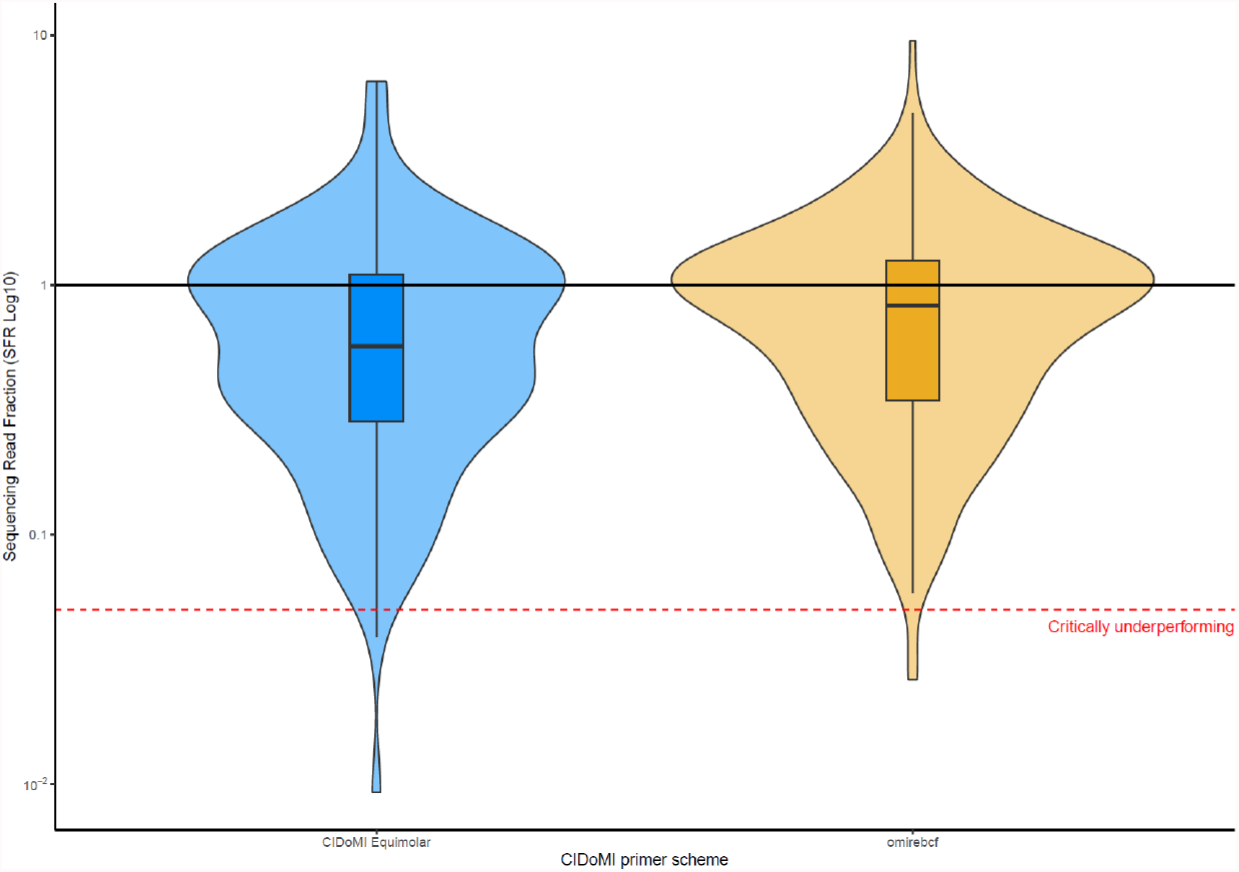
Violin and box plots of sequence read fractions (SRF) of equimolar CIDoMI primers (blue) and rebalanced CIDoMI primers (yellow). Median SRF for the equimolar CIDoMI protocol was 0.83, and rebalanced CIDoMI primers was 0.91.

## References

1. Brito, A.F., et al., Global disparities in SARS-CoV-2 genomic surveillance. 2021, Cold Spring Harbor Laboratory.

2. Koyama, T., D. Platt, and L. Parida, Variant analysis of SARS-CoV-2 genomes. Bull World Health Organ, 2020. 98(7): p. 495–504.

3. ARTICNetwork, SARS-CoV-2 ARTIC v3 Illumina library construction and sequencing protocol V.4. 2020, protocols.io.

4. Freed, N.E., et al., Rapid and inexpensive whole-genome sequencing of SARS-CoV-2 using 1200 bp tiled amplicons and Oxford Nanopore Rapid Barcoding. Biol Methods Protoc, 2020. 5(1): p. bpaa014.

5. Eden, J.S., et al., An emergent clade of SARS-CoV-2 linked to returned travellers from Iran. Virus Evol, 2020. 6(1): p. veaa027.

6. Martin, D.P., et al., Selection analysis identifies unusual clustered mutational changes in Omicron lineage BA.1 that likely impact Spike function. bioRxiv, 2022.

7. Cao, Y., et al., Omicron escapes the majority of existing SARS-CoV-2 neutralizing antibodies. Nature, 2022. 602(7898): p. 657–663.

8. Ma, W., et al., Genomic Perspectives on the Emerging SARS-CoV-2 Omicron Variant. Genomics Proteomics Bioinformatics, 2022.

9. Pru, B.M., Variants of SARS CoV-2: mutations, transmissibility, virulence, drug resistance, and antibody/vaccine sensitivity. Front Biosci (Landmark Ed), 2022. 27(2): p. 65.

10. Cathcart, A.L., et al., The dual function monoclonal antibodies VIR-7831 and VIR-7832 demonstrate potent in vitro and in vivo activity against SARS-CoV-2. bioRxiv 2021.

11. Illingworth, C.J.R., et al., On the effective depth of viral sequence data. Virus Evol, 2017. 3(2): p. vex030.

12. O’Toole, Á., et al., Assignment of Epidemiological Lineages in an Emerging Pandemic Using the Pangolin Tool. Virus Evolution, 2021.

13. Rambaut, A., et al., A dynamic nomenclature proposal for SARS-CoV-2 lineages to assist genomic epidemiology. Nature Microbiology, 2020. 5(11): p. 1403–1407.

14. Basile, K., et al., Cell-based Culture Informs Infectivity and Safe De-Isolation Assessments in Patients with Coronavirus Disease 2019. Clin Infect Dis, 2020.

15. Basile, K., et al., Cell-based culture of SARS-CoV-2 informs infectivity and safe de-isolation assessments during COVID-19. Clin Infect Dis, 2020.

16. Lam, C., et al., SARS-CoV-2 Genome Sequencing Methods Differ In Their Ability To Detect Variants From Low Viral Load Samples. J Clin Microbiol, 2021.

17. Rockett, R.J., et al., Co-infection with SARS-CoV-2 Omicron and Delta variants revealed by genomic surveillance. Nat Commun, 2022. 13(1): p. 2745.

18. R&D, D.P., et al., COVID-19 ARTIC v4.1 Illumina library construction and sequencing protocol - tailed method V.2. 2022, protocols.io.

19. Quick, J., et al., Multiplex PCR method for MinION and Illumina sequencing of Zika and other virus genomes directly from clinical samples. Nat Protoc, 2017. 12(6): p. 1261–1276.

20. Rockett, R.J., et al., Revealing COVID-19 transmission in Australia by SARS-CoV-2 genome sequencing and agent-based modeling. Nat Med, 2020. 26(9): p. 1398–1404.

21. Islam, M.R., The SARS-CoV-2 Omicron (B.1.1.529) variant and the re-emergence of COVID-19 in Europe: An alarm for Bangladesh. Health Science Reports, 2022. 5(2): p. e545.

22. Khandia, R., et al., Emergence of SARS-CoV-2 Omicron (B.1.1.529) variant, salient features, high global health concerns and strategies to counter it amid ongoing COVID-19 pandemic. Environ Res, 2022. 209: p. 112816.

23. ARTICNetwork, SARS-CoV-2 V4.1 update for Omicron variant. 2021.

24. Viana, R., et al., Rapid epidemic expansion of the SARS-CoV-2 Omicron variant in southern Africa. Nature, 2022. 603(7902): p. 679–686.

25. Omicron variant in confirmed NSW cases. 2021, NSW Health.

26. Nadeau, S.A., et al., Swiss public health measures associated with reduced SARS-CoV-2 transmission using genome data. Sci Transl Med, 2023. 15(680): p. eabn7979.

27. Nielsen, B.F., et al., Host heterogeneity and epistasis explain punctuated evolution of SARS-CoV-2. PLoS Comput Biol, 2023. 19(2): p. e1010896.

28. Harvey, W.T., et al., SARS-CoV-2 variants, spike mutations and immune escape. Nat Rev Microbiol, 2021. 19(7): p. 409–424.

29. Corey, L., et al., SARS-CoV-2 Variants in Patients with Immunosuppression. N Engl J Med, 2021. 385(6): p. 562–566.

30. Lafzi, A., et al., Tutorial: guidelines for the experimental design of single-cell RNA sequencing studies. Nat Protoc, 2018. 13(12): p. 2742–2757.

